# Evolution of polyamine resistance in *Staphylococcus aureus* through modulation of potassium transport

**DOI:** 10.1101/2024.06.15.599172

**Authors:** Killian Campbell, Caitlin H. Kowalski, Kristin M. Kohler, Matthew F. Barber

## Abstract

Microbes must adapt to diverse biotic and abiotic factors encountered in host environments. Polyamines are an abundant class of aliphatic molecules that play essential roles in fundamental cellular processes across the tree of life. Surprisingly, the bacterial pathogen *Staphylococcus aureus* is highly sensitive to polyamines encountered during infection, and acquisition of a polyamine resistance locus has been implicated in spread of the prominent USA300 methicillin-resistant *S. aureus* lineage. At present, alternative pathways of polyamine resistance in staphylococci are largely unknown. Here we applied experimental evolution to identify novel mechanisms and consequences of *S. aureus* adaption when exposed to increasing concentrations of the polyamine spermine. Evolved populations of *S. aureus* exhibited striking evidence of parallel adaptation, accumulating independent mutations in the potassium transporter genes *ktrA* and *ktrD*. Mutations in either *ktrA* or *ktrD* are sufficient to confer polyamine resistance and function in an additive manner. Moreover, we find that ktr mutations provide increased resistance to multiple classes of unrelated cationic antibiotics, suggesting a common mechanism of resistance. Consistent with this hypothesis, ktr mutants exhibit alterations in cell surface charge indicative of reduced affinity and uptake of cationic molecules. Finally, we observe that laboratory-evolved ktr mutations are also present in diverse natural *S. aureus* isolates, suggesting these mutations may contribute to antimicrobial resistance during human infections. Collectively this study identifies a new role for potassium transport in *S. aureus* polyamine resistance with consequences for susceptibility to both host-derived and clinically-used antimicrobials.

## Introduction

*Staphylococcus aureus* is a gram-positive bacterium that colonizes approximately 30% of the human population as well as human-associated animals (1, 2). *S. aureus* is most frequently isolated from the anterior nares, where it is considered a commensal member of the human microbiota (3, 4). However, *S. aureus* is also capable of colonizing the skin and is a leading cause of soft skin and tissue infections (SSTIs) as well as other invasive conditions including bloodstream infections, pneumonia, osteomyelitis, and endocarditis (4–8). *S. aureus* is a frequent agent of antibiotic-resistant infections, with methicillin resistant *S. aureus* (MRSA) causing between 10,000 and 20,000 deaths in the United States annually (8, 9). Recent global estimates indicate that *S. aureus* is a leading cause of both total bacterial infections and antibiotic resistant infections in humans across geographic and socioeconomic boundaries (10, 11).

Previous studies have identified a number of biotic and abiotic factors that limit *S. aureus* colonization in distinct host niches (12–15). In particular, polyamines are an abundant class of aliphatic cationic molecules known to restrict *S. aureus* growth during infection (16, 17). Polyamines are characterized by the presence of at least two primary amino groups and include putrescine, agmatine, spermidine, and spermine (18, 19). Polyamines are byproducts of arginine metabolism and may be synthesized by different series of enzymatic steps depending on the organism (20). Polyamine metabolism and functions are well-studied in eukaryotes, where they play pleiotropic roles in normal cellular physiology (21–23). Due to their net positive charge, polyamines are capable of binding to ribosomes and DNA during protein production and DNA replication, respectively (18, 24). In mammals, the enzymes that produce and breakdown polyamines are controlled at multiple transcriptional levels, resulting in a tightly-regulated pool of polyamines within a cell at any given time (19, 25).

Given the numerous roles that polyamines play in essential cellular functions, polyamine synthesis was once believed to be conserved across the entire tree of life (26). This dogma was overturned by previous work demonstrating that multiple bacterial genera have lost the ability to synthesize polyamines *de novo* (21). Even among bacteria that have lost the ability to synthesize polyamines, supplementing media with exogenous polyamines often results in modest growth improvements (27, 28). In marked contrast to other bacteria, polyamines are highly bactericidal to nearly all *Staphylococcus* species surveyed with particularly potent effects on *S. aureus* (16). A key exception are *S. aureus* strains within the USA300 lineage, which exhibit enhanced polyamine resistance (28, 29). USA300 is a lineage of community-acquired MRSA (CA-MRSA) that emerged in the early 2000s. CA-MRSA strains differ from previously dominant hospital-acquired (HA) MRSA strains both in disease pathology and the fact that they are not limited to nosocomial settings (30). CA-MRSA infections began to rise in the early 1990s and were predominantly caused by members of the USA400 and USA100 lineages (30). Subsequent emergence of USA300 CA-MRSA strains led to the rapid replacement of other CA-MRSA lineages to become the dominant agents of *S. aureus* SSTIs in North America (31). Given the global spread of USA300, it was hypothesized that this lineage is both hyper-transmissible and hyper-virulent, contributing to its impressive evolutionary success (30). Subsequent work revealed that USA300 isolates almost uniformly encode genes for the Panton-Valentine leucocidin as well as mutations in the global transcription regulators *sae* and *agr* which lead to increased expression of toxins, contributing to enhanced virulence (30–33). However, these traits are shared amongst other MRSA lineages, and it wasn’t until the discovery of the arginine-catabolic mobile element (ACME) unique to USA300 that a new genetic determinant of fitness was identified in this group (28, 29, 34). ACME is an approximately 31kb genomic island that contains at least 33 putative genes (28, 29). Evidence suggests that ACME was assembled in *Staphylococcus epidermidis* and horizontally transferred to the most recent common ancestor of USA300 (28). Notably, ACME contains a spermine acetyltransferase gene, *speG*, which confers resistance to polyamines that *S. aureus* encounters in the infection environment (16, 28).

The ACME locus contains several genes related to arginine metabolism, which produce ammonia allowing these strains to efficiently colonize the acidic environment of the skin (29). As a result, ACME promotes efficient colonization of skin and promotes infection success after colonization (16, 28, 35). At present, the basis of polyamine toxicity and mechanisms by which non-USA300 *S. aureus* may adapt to these molecules are largely unknown. While ACME and *speG* in particular provide an example of how polyamine resistance may evolve, alternative genetic or molecular mechanisms of resistance have not been characterized. To address this question, we experimentally evolved populations of *S. aureus* under increasing concentrations of spermine and surveyed the consequences of resistance evolution across three different strain backgrounds. Our findings identify both new genes and new mechanisms conferring polyamine resistance in this major bacterial pathogen.

## Results

### Experimental evolution of spermine resistance in S. aureus

To identify mechanisms of polyamine resistance in *S. aureus*, we performed a serial-passaging experiment in which populations of *S. aureus* were continuously exposed to increasing concentrations of spermine (Figure 1A). Initially, single colonies of *S. aureus* strains MN8, HFH-30364, and RN4850 were revived from glycerol stocks on tryptic soy agar (TSA) plates. Both *S. aureus* MN8 and RN4850 are methicillin sensitive strains while HFH-30364 is an MRSA strain from the USA400 lineage (Table 1). Replicate populations of each strain were then inoculated in tryptic soy broth media (TSB) to perform the passaging experiment. Spermine was selected as a representative polyamine as it has a well-documented bactericidal effect on *S.* aureus (16) and is abundant during the post inflammatory phase of wound healing in murine models (17). Each day, a 1/100 aliquot of each replicate culture was transferred to fresh TSB media with a pre-determined concentration of spermine (or no spermine for the control populations). Populations were initially exposed to sub-inhibitory conditions of spermine (2 mM) and passaged daily, for 10 days, until reaching a final concentration of 7 mM spermine. Genomic DNA was also collected to perform whole-population sequencing of MN8 samples taken throughout the evolution experiment. For RN4850 and HFH-30364, we conducted whole-genome, population-level sequencing upon completion of the passaging experiment (Fig. 1A).

**Figure 1:**
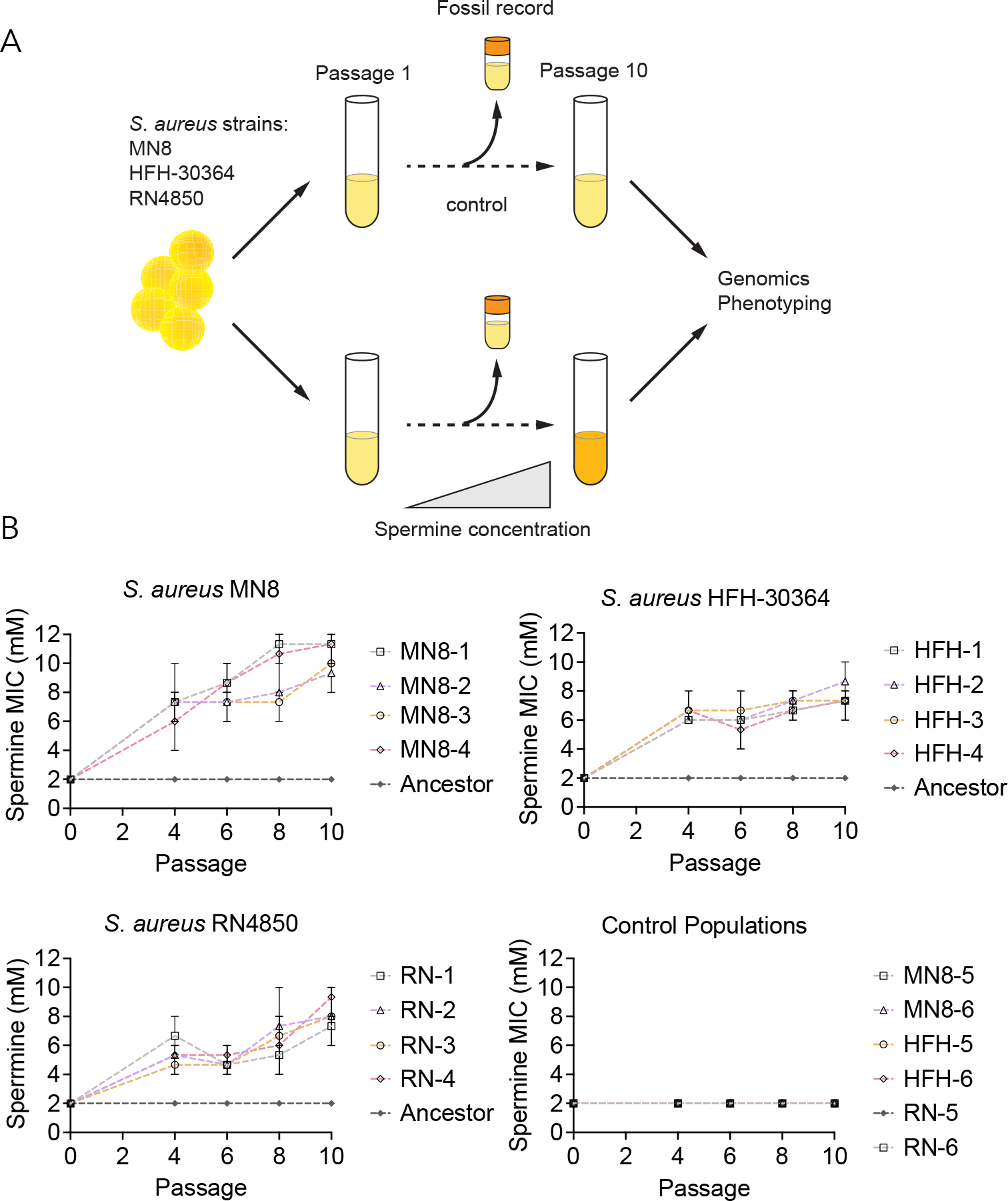
Experimental evolution of spermine resistance in *Staphylococcus aureus*. **A.** Experimental evolution overview. Populations of each strain of *S. aureus* (MN8, HFH-30364, and RN4850) were revived from glycerol stocks on to tryptic soy agar (TSA). For each strain background, four single colonies were picked to establish replicate populations to undergo exposure to spermine and two colonies were picked to establish replicate control lines to be passaged in TSB alone. Populations were passaged at a fixed dilution daily, and spermine-exposed lines were continuously exposed to spermine. Glycerol stocks containing aliquots of the evolving populations were saved periodically as a “fossil record.” At the end of the experiment, glycerol stocks were revived to perform phenotypic analyses and to isolate DNA for sequencing. **B.** Spermine minimal inhibitory concentration (MIC) was calculated over the course of the evolution experiment. Glycerol stocks from throughout the experiment were revived on TSA plates, then single colonies were picked and spermine MIC was measured. Shapes and error bars represent average and range of MIC values measured. Each point represent the average of a separate experiment conducted in triplicate.

**Table 1.** Mutations observed during experimental evolution. A subset of mutations that arose in each lineage of each strain background during the evolution experiment. Mutations reported were detected using breseq from while genome sequencing from DNA isolated at passage 10 of the evolution experiment.

| Population | *ktr* potassium transport complex |  | ATP-synthase machinery |  |  |  | *pgl* |
| --- | --- | --- | --- | --- | --- | --- | --- |
|  | *ktrA* | *ktrD* | *atpG* | ATP synthase alpha subunit | ATP synthase delta subunit | ATP synthase gamma subunit |  |
| MN8-1 | E76G | Q450K, S135L |  |  |  |  |  |
| MN8-2 |  |  |  |  |  |  |  |
| MN8-3 | E76G | M391I |  |  |  |  |  |
| MN8-4 | A178V | G94C |  |  |  |  |  |
| HFH-1 |  |  |  | P95L |  |  | G4E, H306Y |
| HFH-2 | E151V |  | T242P |  |  |  | G108V |
| HFH-3 |  |  |  |  |  |  | G278V, G323D |
| HFH-4 |  |  |  |  |  |  | G120S, R318S |
| RN-1 |  |  |  |  |  |  | R245V, T10P |
| RN-2 |  |  |  |  |  | Δ69 bp | P234L, S41F |
| RN-3 |  |  |  |  |  | L261* | A245V |
| RN-4 |  |  |  |  | Q170* |  |  |

We observed that all populations in each of the three strain backgrounds evolved increased spermine resistance within 10 passages of laboratory evolution. Specifically, both *S. aureus* MN8 and HFH-30364 populations evolved levels of resistance at least 4-fold greater than the minimal inhibitory concentration (MIC) of the ancestral strain (Figure 1B). *S. aureus* RN4850 exhibited a slightly more variable resistance profile, with most replicate populations evolving an MIC nearly four times the ancestral levels (Figure 1B). Individual colonies tested from each population displayed slight differences in resistance profiles, likely reflecting heterogeneity in mutations carried by individual clones. *S. aureus* MN8 displayed the largest fold-change in spermine MIC, with the MIC of individual colonies measuring between 5-6X the ancestral MIC levels (Figure 1B). We noted that the dynamics of resistance evolution as measured by spermine MIC varied between strain backgrounds. *S. aureus* MN8 steadily evolved increased resistance throughout the passaging experiment, while HFH-30364 experienced a single, large increase in resistance change between passages 0-4, and RN4850 experienced two large increases in resistance from passage 0-4 and passage 8-10 (Figure 1B). Together this experiment revealed that *S. aureus* can readily evolve resistance to spermine, while the dynamics of resistance evolution differ between strains (Figure 1B). Given that *S. aureus* MN8 populations evolved the highest magnitude change of resistance relative to the ancestor, we chose to focus on characterizing the genetic basis of spermine resistance in this strain background for future experiments.

### Genetic determinants of spermine resistance

We next sought to determine the genetic basis of polyamine resistance in our evolved *S. aureus* populations. Whole-population DNA samples were extracted from each replicate population throughout the experiment (MN8) and at the end of the passaging experiment (HFH-30364, RN4850). Additionally, we extracted DNA from ancestral clones to generate a reference sequence for variant calling. *S. aureus* MN8 populations exhibited striking evidence of convergent evolution, with three of the four spermine-exposed populations harboring mutations in different components of the *ktr* potassium transport complex (Table 1). The Ktr system has been characterized in *Bacillus subtilis* as a constitutively expressed moderate-affinity potassium transporting system, and has only recently been described in *S. aureus* (36, 37). This complex is composed of two proteins: KtrA, which forms a regulatory octameric ring subunit in the cytoplasm, and KtrD, a dimeric transmembrane channel that facilitates potassium import. Previous work has implicated the Ktr complex in both osmotic and alkaline stress tolerance (36–38). The Ktr complex also contributes to survival to a variety of antibacterial agents under low potassium conditions, as Δ*ktrA* deletion mutants have increased sensitivity to aminoglycoside antibiotics (36, 37). Despite this, no study to date has directly linked Ktr complex function and polyamine resistance.

MN8-derived populations exhibited the highest levels of polyamine resistance of the three strain backgrounds tested and were the only populations carrying high-frequency mutations in *ktr* genes (Table 1). We were therefore motivated to further investigate the evolutionary dynamics of spermine resistance in the MN8 populations. We leveraged whole-genome, whole-population sequencing to track the frequencies of *ktr* mutations over time. In each of the three replicate populations containing *ktr* mutations (MN8-1, MN8-3, MN8-4), nonsynonymous mutations occurred in both *ktrA* and *ktrD* at high frequencies by the final passage (Figure 2A-C). Specifically, a single nonsynonymous mutation in *ktrA* had fixed by passage 10 in replicate populations MN8-1 and MN8-4, and had nearly fixed in MN8-3 (84% allele frequency) (Figure 2A, 2C).

**Figure 2.**
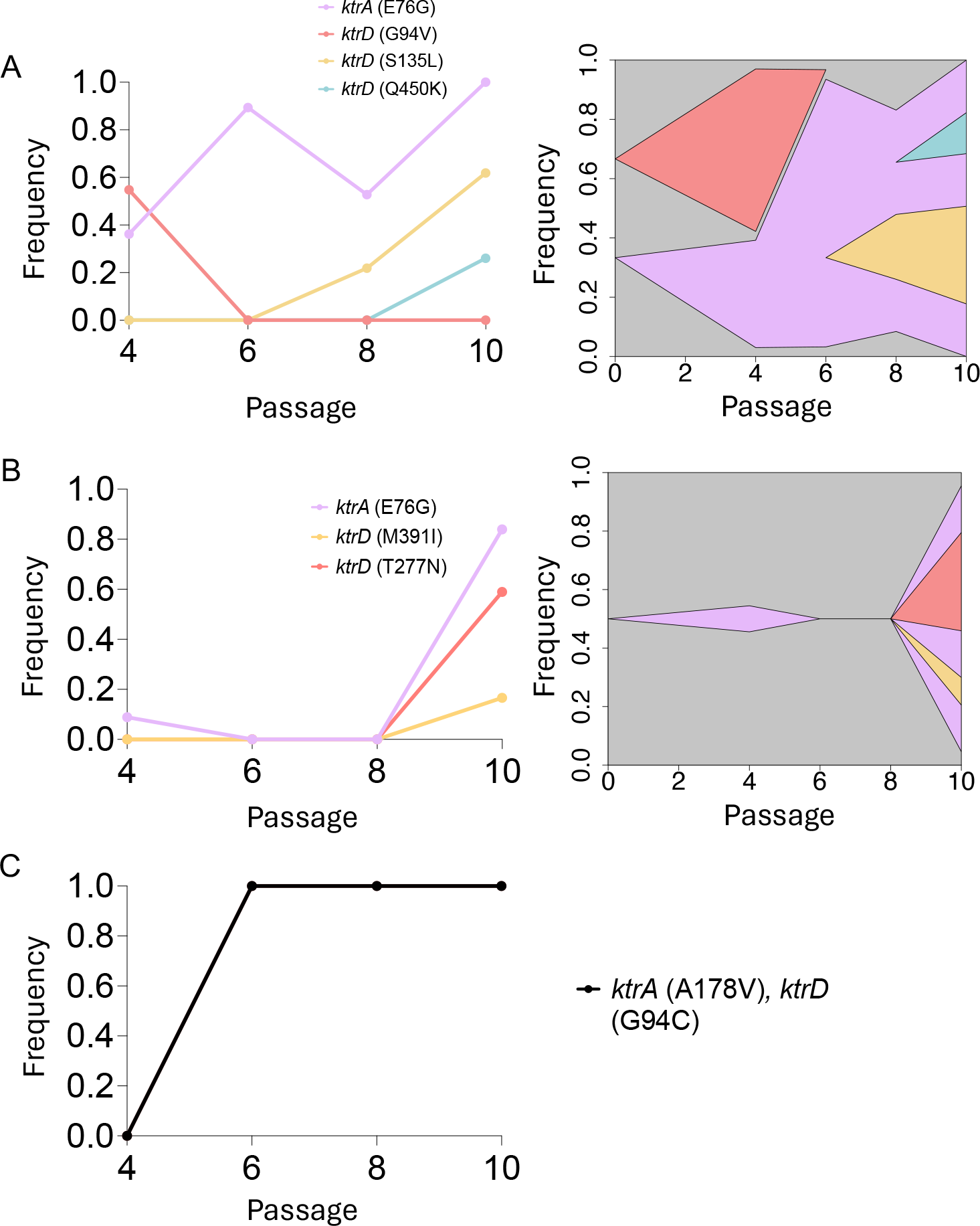
Allele frequency dynamics and linkage of *ktr* mutations over the course of experimental evolution. Allele frequencies and linkage were determined for ktrA and ktrD mutations in the MN8-1 (**A**), MN8-3 (**B**), and MN8-4 (**C**) populations. For all allele frequency line plots, breseq was run in polymorphism mode to determine the frequency of *ktrA* and *ktrD* single nucleotide polymorphisms (SNPs) detected in the population throughout the evolution experiment. To determine the linkage of *ktr* alleles present in each population, a combination of PCR and Sanger sequencing was used. For both MN8-1 and MN8-3, 16 different colonies were picked and colony PCR was conducted on both the *ktrA* and *ktrD* locus for each colony. PCR products were sent for Sanger sequencing, and sequences were aligned to detect which SNPs were present in each clone. Sanger sequencing revealed that clones in both MN8-1 and MN8-3 could contain a single SNP in *ktrD* on the background of a single snp in *ktrA*.

Notably, only a *single ktrA* allele was maintained in in any of the populations by the final passage, whereas multiple *ktrD* alleles co-occurred in both MN8-1 and MN8-3 (Figure 2A, 2B). The allele frequency dynamics were also different throughout each of the replicate populations. Single mutations in *ktrA* and *ktrD* arose early and fixed in MN8-4 by passage 6 (Figure 2C). However, the dynamics for MN8-1 and MN8-3 were more varied. In MN8-1, a single mutation in *ktrA* largely persisted until generation 8, when multiple *ktrD* mutations were detected in the population at low frequencies (Figure 2A. 2B). MN8-3 displayed similar characteristics to MN8-4, in which all *ktr* mutations arose late in the experiment between passages 8 and 10 (Figure 2B, 2C). We next picked multiple clones from the terminal passage in both MN8-1 and MN8-3 to determine the linkage of these *ktr* mutations. While multiple mutations were present in the *ktrD* gene in MN8-1, they existed in different lineages, and always in combination with the *ktrA* mutation (Figure 2A). This was also true in MN8-3, where a single mutation in *ktrD* was found in combination with a single mutation in *ktrA* (Figure 2B). Despite multiple *ktrD* mutations existing at the population level, we did not detect any lineages within MN8-3 carrying multiple *ktrD* mutations (Figure 2B). Our findings suggest that resistance to spermine was largely conferred through a combination of single mutations in both *ktrA* and *ktrD*.

### Single amino acid changes in ktr genes confer additive polyamine resistance

To determine whether single mutations in *ktr* genes are sufficient to confer spermine resistance in *S. aureus*, we generated targeted substitutions in both KtrA and KtrD by homologous recombination in strain MN8. Specifically, the histidine at position 47 of KtrA was substituted for a tyrosine (H47Y), and the glycine at position 94 of KtrD was substituted for a valine (G94V) (Figure 3A). These mutations were recovered in a previously conducted pilot evolution experiment (Supplemental Figure 4) and are in proximity to mutations recovered in the evolution experiment described above. We then measured spermine MIC of each single mutant, as well as the *ktrA/ktrD* double mutant. We found that each single mutation conferred partial resistance to spermine, with an MIC of 6-8 mM for both the KtrA H47Y and KtrD G94V alleles (Figure 3B). However, the *ktrA/ktrD* double mutant conferred a similar level of resistance to spermine that we observed at the end of the evolution experiment (7 mM), making this combination of mutations sufficient to provide the level of resistance observed in these populations (Figure 3B). Together these results demonstrate that evolved mutations in *ktrA* and *ktrD* are sufficient to confer polyamine resistance in *S. aureus* in an additive manner. We also confirmed the ability of these strains to survive spermine concentrations by enumerating colony forming units per milliliter of culture (CFU/mL) (Figure 3B). We observed a ∼100-fold reduction in recovered CFUs for the ancestral strain when incubated for 24 hours in 4 mM spermine compared to the no spermine control. In contrast, there was no reduction in CFUs observed for either of the single mutants in 4 mM versus 0 mM spermine (Figure 3B). We similarly did not observe a significant reduction in CFU/mL for the *ktrA/ktrD* double mutant when incubated with 4mM or 8 mM spermine, reflecting the ability of single amino acid changes to confer spermine resistance (Figure 3B).

**Figure 3.**
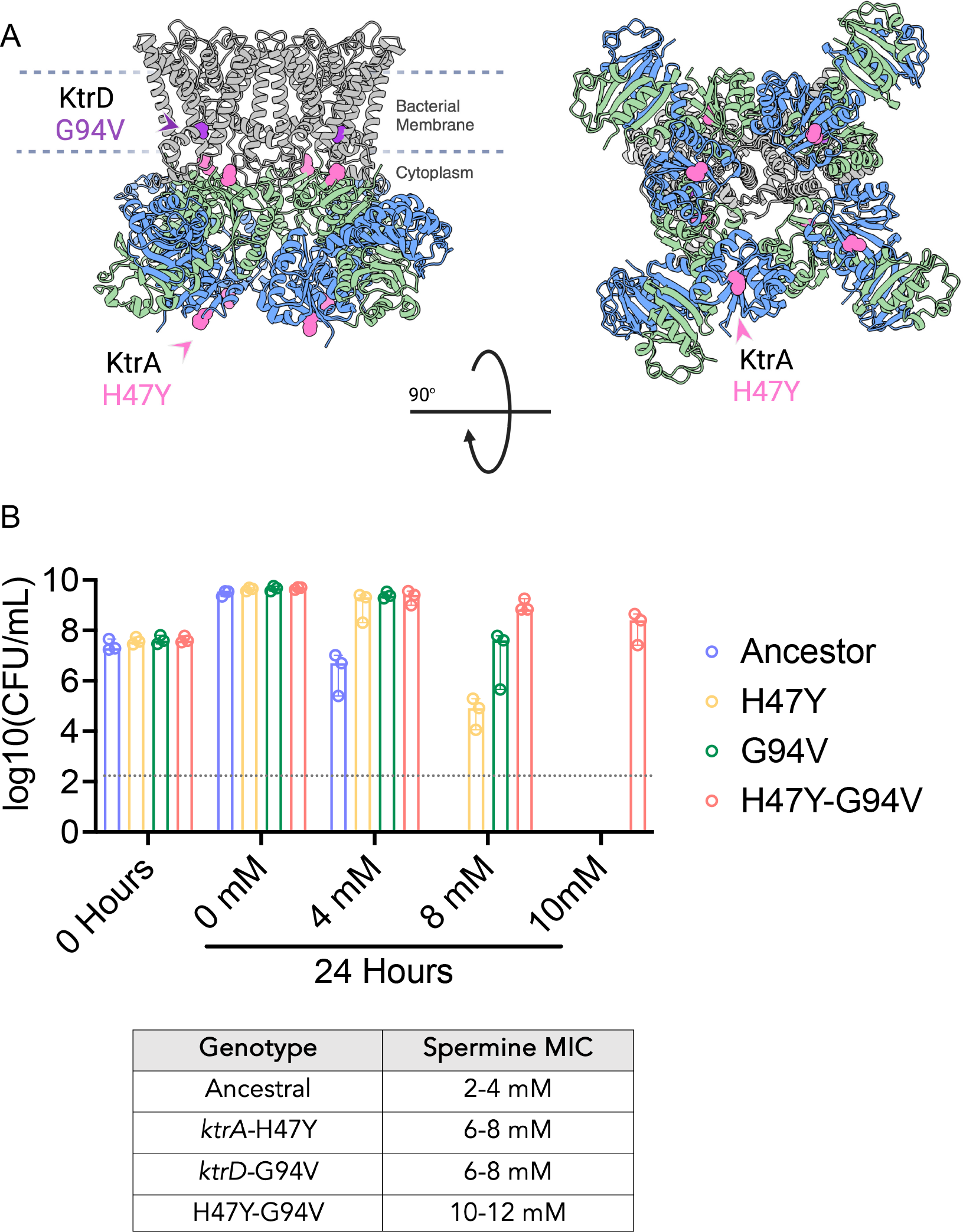
Evolved mutations in the KtrAD complex confer additive spermine resistance in S. aureus. **A.** Predicted structure of the KtrAD complex generated using ChimeraX. *S. aureus ktrA* and *ktrD* gene sequences were used to model the KtrAD complex based on the published KtrAD crystal structure from *Bacillus subtilis* (PDB ID: 4J7C). Mutations introduced into the MN8 strain background are highlighted in pink (KtrA) and purple (KtrD). **B.** Colony forming units (CFUs) recovered from MN8 *S. aureus* strains incubated in different concentrations of spermine for 24 hours (top), with corresponding spermine MIC values listed in the table (bottom). Gray dotted line represents limit of detection.

### Evolved ktr mutations alter S. aureus antibiotic susceptibility

Previous reports indicate that the *ktr* potassium transport system plays an important role in antimicrobial resistance and alkaline stress tolerance in *S. aureus* (37). We therefore sought to determine if our evolved *ktr* mutations also altered the ability of *S. aureus* to tolerate other stressors beyond spermine. We measured the change in MIC for a panel of antibiotics in the ancestral strain MN8 as well as the engineered *ktrA* (H47Y) and *ktrD* (G94V) mutants (Figure 4A). Previous findings revealed that *ktr* null mutants sensitize *S. aureus* to aminoglycoside antibiotics (36). In contrast to null mutants, the evolved *ktrA/ktrD* double-mutant exhibits an approximately 12-fold increase in aminoglycoside MIC compared to the ancestor (Figure 4A). Aminoglycoside antibiotics rely upon an intact proton motive force to be internalized across the *S. aureus* cell membrane, where they can interfere with protein translation (39, 40). We therefore hypothesized that the *ktr* mutants identified in this study function by altering membrane potential and preventing aminoglycoside internalization. Potassium transport, and *ktr-*mediated potassium transport in particular, is known to regulate membrane potential (36, 41, 42). We detected a modestly depolarized membrane potential in the *ktr* double mutants compared to the ancestral strain (Figure 4B). Despite this small but reproducible difference in membrane potential, we were not convinced that this was the only factor contributing to antimicrobial resistance in the *ktr* mutants. Prior to internalization into the bacterial cell, uptake is governed by electrostatic attraction between the positively-charged aminoglycoside molecule and the negatively-charge bacterial cell wall (43). Given the polycationic nature of both spermine and aminoglycosides at physiological pH and the observed resistance of the *ktr* mutants to these molecules, we hypothesized that an alteration in cell-surface charge, coupled with membrane depolarization, could explain the observed change in resistance. To test this hypothesis, we measured survival of the *ktr* double mutant in polymyxin B, an unrelated polycationic antibiotic that has a +5-charge at physiological pH (44). Consistent with our hypothesis, the *ktr* double mutant survived significantly better than the ancestor in the presence of 1 mg/mL polymyxin B (Fig 4C). To assess changes in bacterial cell surface charge, we measured cytochrome C binding in the ancestral and *ktr* double mutant strains (Figure 4D). Cytochrome C binding is a convenient proxy to infer bacterial cell-surface charge as it is highly cationic and binds effectively to the negatively charged bacterial cell wall (45). We detected a significant increase in the percent of unbound cytochrome C in the *ktr* mutant when compared to the ancestral strain, consistent with an increase in cell surface charge conferred by *ktr* mutations (Figure 4D). Taken together, our findings suggest a novel mechanism by which modulation of potassium transport complex function confers resistance to cationic antimicrobial compounds through alteration of cell surface charge and possibly membrane potential.

**Figure 4.**
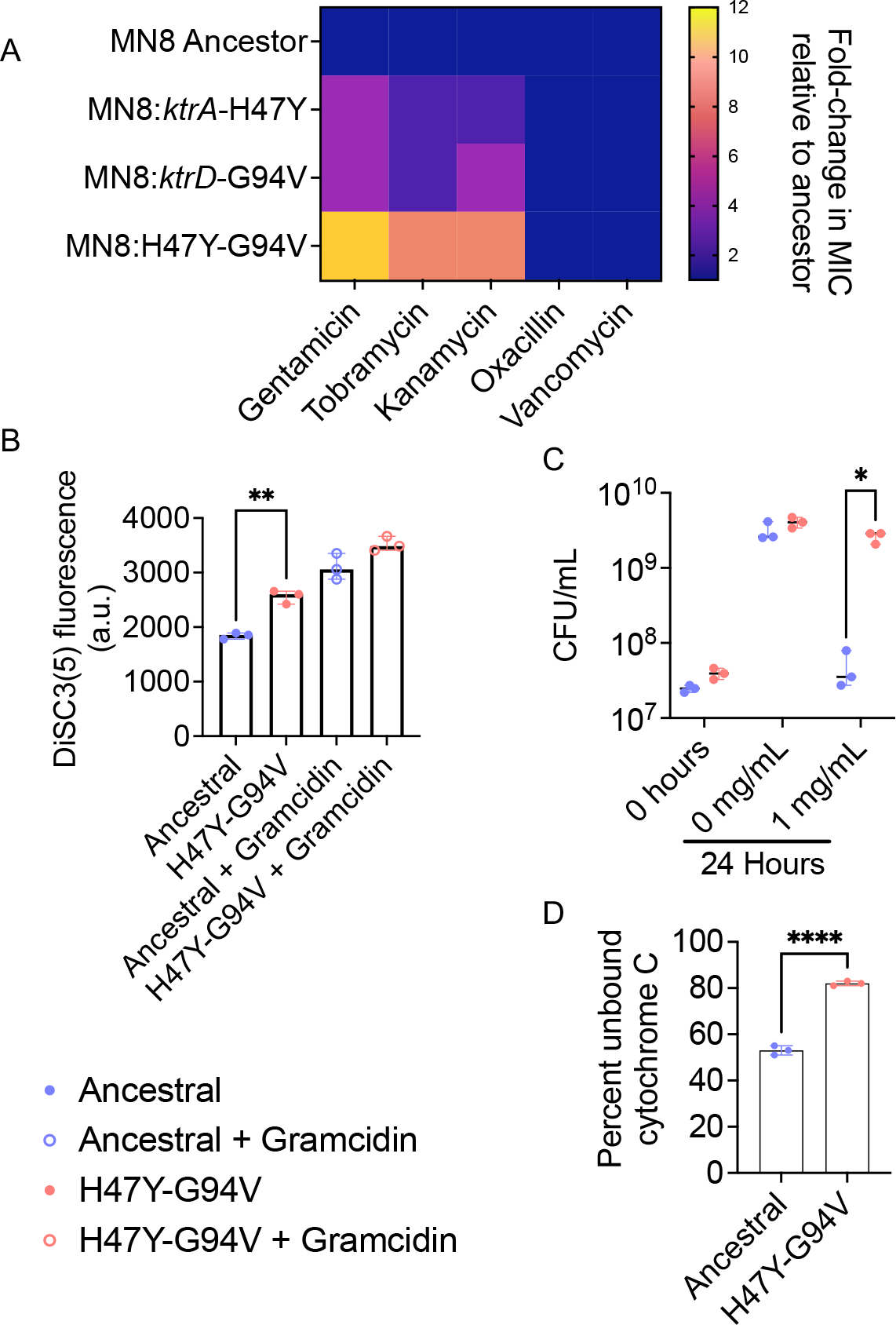
KtrAD complex mutations alter antibiotic resistance, membrane potential, and cell-surface charge. **A.** Heatmap depicting fold-change in MIC for indicated antibiotics in *S. aureus* mutant strains relative to the ancestor. Values represent the average of three experiments each conducted in triplicate. **B.** 3,3’-Dipropylthiadicarbocyanine Iodide (DiSC3(5)) fluorescence of different engineered strains. Fluorescence readings are normalized to OD600 measurements for each respective strains. Points on the graph each represent average of a different experiment conducted in triplicate. Asterisks represent significant difference (P<0.01) as detected by an unpaired Student’s T-test. **C.** CFUs of the ancestral MN8 strain or double *ktrAD* mutant measured after 24 hours of incubation with polymyxin B. Points represent average of a different experiment conducted in triplicate. Asterisks represent significant difference (P<0.05) in means between two groups compared as detected by an unpaired T-test. **D.** Percent unbound cytochrome C measured via absorbance at 410 nm in a spectrophotometric plate reader. Percent unbound cytochrome C was calculated relative to the absorbance of cytochrome C alone in buffer without cells. Asterisks represent significant difference (P<0.001) as determined by an unpaired T-test.

### Exogenous cations reduce spermine and aminoglycoside toxicity in S. aureus

Given that *ktr* mutants identified in this study exhibit opposite drug resistance phenotypes compared to published *ktr* deletion mutants (37), we hypothesized that our evolved mutants function by increasing potassium import relative to wild-type (WT) cells. To test if elevated levels of potassium can similarly protect against spermine and aminoglycoside toxicity, we generated a chemically defined medium lacking excess potassium (38). This allowed us to supplement the growth medium with defined amounts of potassium in the form of KCl. Supplementation with 250 mM KCl significantly increased *S. aureus* survival in the presence of 4 mM spermine, consistent with our hypothesis (Fig. 5A). In addition, we recovered over 10^5^ CFUs when *S. aureus* MN8 was exposed to 6 mM spermine for 24 hours in 250 mM KCl, whereas the control condition (10 mM KCl) yielded no detectable CFUs (Fig. 5A). Similar to spermine, we observed reduced sensitivity of *S. aureus* to all of the aminoglycoside antibiotics tested (measured as an increase in MIC) with 250 mM KCl compared to the 10 mM KCl control (Fig 5C). We next considered whether this protective effect was specific to potassium, or generalizable to other monovalent cations. To address this question, we repeated the experiments described above supplementing with NaCl instead of KCl. We found that the addition of NaCl at the same concentrations conferred a similar level of protection to KCl against spermine and aminoglycosides (Figure 5B, 5C). Together these findings demonstrate that, like genetic alterations in potassium transport function, excess levels of exogenous cations are sufficient to enhance polyamine resistance in *S. aureus*.

**Figure 5.**
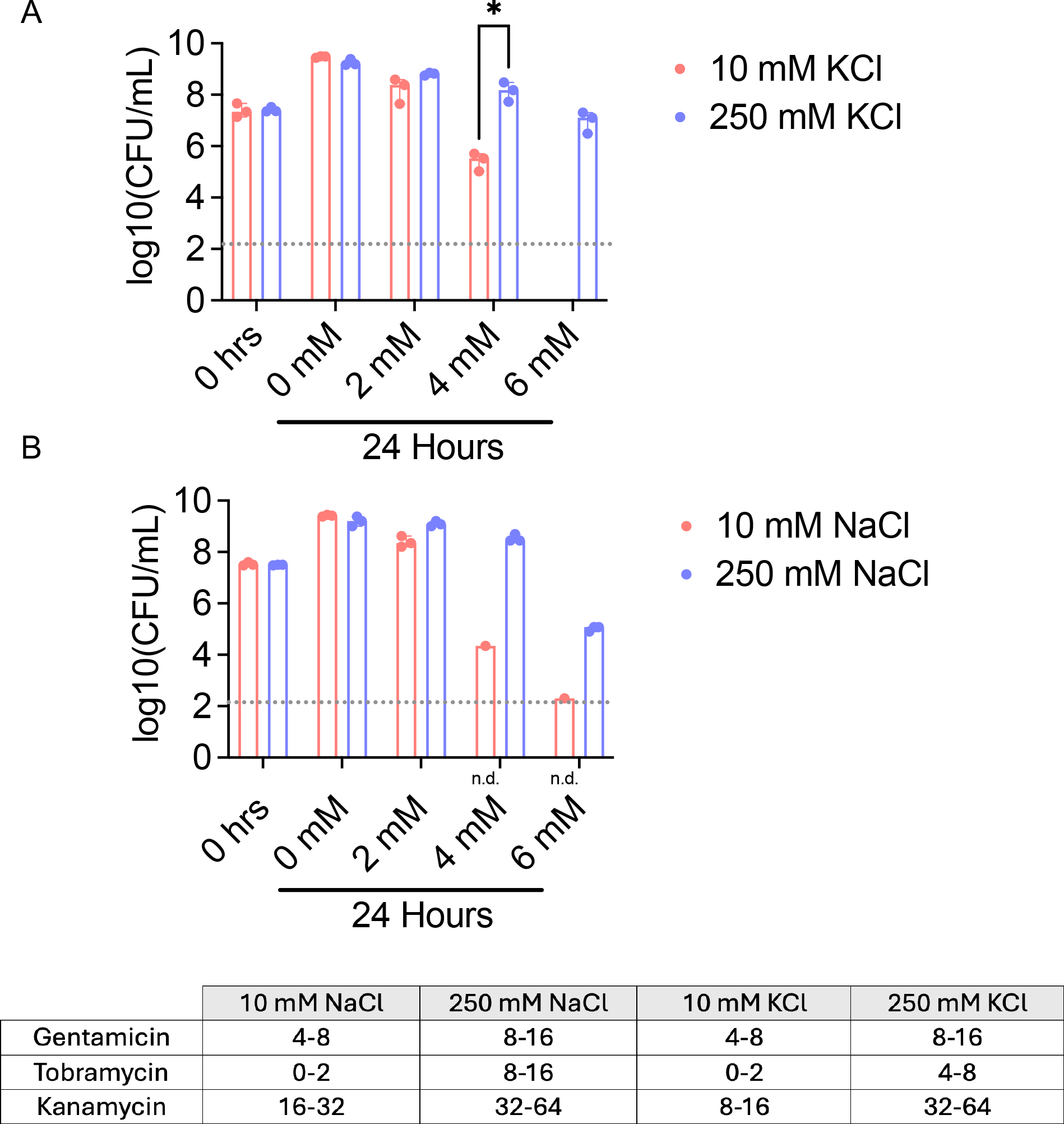
Exogenous potassium or sodium is sufficient to ameliorate spermine toxicity in S. aureus. **A.** Enumeration of CFUs after incubation of S. aureus strain MN8 with spermine in the presence of exogenous KCl. Gray dotted line denotes limit of detection. Asterisks denote significant difference in means between two treatments as detected by an unpaired t-test (P<0.05). **B.** Enumeration of CFUs after incubation of S. aureus strain MN8 with spermine in the presence of exogenous NaCl. Gray dotted line denotes limit of detection. N.d. denotes no recoverable CFUs detected for 2/3 replicates in those conditions. Table depicting MIC values in microgram/mL for different aminoglycoside antibiotics incubated for 24 hours in different amounts of KCl or NaCl.

## Discussion

Our study identifies a previously unknown role for the Ktr potassium transport complex in polyamine resistance in *S. aureus*. A recent study also recovered mutations in *ktr* genes after passaging *S. aureus* in the presence of a synthetic polyamine, although the effects of these mutations were not reported (46). It is notable that phenotypes conferred by single amino acid changes in the Ktr complex recovered during experimental evolution are markedly different from Δ*ktr* strains described previously (36). Evolved *ktrA* and *ktrD* mutations uniformly decreased sensitivity to aminoglycoside antibiotics and spermine, suggesting a gain of function phenotype. Our experiments supplementing the media with cations further supports this, as abundant levels of exogenous cations largely protect from the toxic effects of both spermine and aminoglycoside antibiotics (Fig. 5). In contrast, Δ*ktr* strains are highly sensitive to aminoglycoside antibiotics relative to WT (36). These same mutations in *ktr* genes altered membrane potential and cell-surface charge, each of which contribute to spermine and aminoglycoside resistance. Links between potassium transport and membrane potential are well-documented, although there is a lack of evidence in the literature regarding the role potassium transport in regulating bacterial cell surface charge (47). Notably, a previous study also found that potassium transport can alter zeta potential (the voltage field that arises from a cell surface) in a red blood cell model (48). Work is ongoing to clarify the connections between Ktr complex function, bacterial physiology, and resistance to antimicrobial agents. Notably, the relationship between cations, polyamines, aminoglycosides, and gram-positive bacteria has been noted before (54). This work demonstrated that aminoglycosides, spermidine, and magnesium cations compete for binding wall-teichoic acids on gram-positive bacterial cell walls. Future studies could aid in further clarifying how alterations in cell-wall architecture influence susceptibility to polyamines and other cationic antimicrobials.

The observation that natural *S. aureus* isolates carry identical mutations in Ktr complex genes as those recovered in our experiments suggests that changes in potassium transport may confer important fitness advantages during human colonization or infection (Supplemental table 2). Notably, we find that these mutations provide enhanced resistance to diverse cationic antimicrobials, potentially through alterations in bacterial cell surface charge (Fig. 4). In *S. aureus,* changes in cell-surface charge can reduce susceptibility to both daptomycin and vancomycin, two antibiotics of choice for treating MRSA infection (49, 50). Additionally, cell-surface charge alterations can promote defense against host-encoded antimicrobial peptides, suggesting a potential strategy for *S. aureus* to establish an infection in the host environment (51). Further studies to understand the contribution of Ktr complex function during human infections or drug resistance could aid in further addressing these questions. In addition, our findings illustrate how combining laboratory experimental evolution with natural strain population genomics can yield useful insights on the genetic basis of pathogen adaptation. In this study, mutations in *ktr* were largely only detected in the MN8 strain background (Table 2). Conversely, mutations in *pgl,* a gene that encodes for a lactonase and acts in the pentose phosphate pathway were not recovered in MN8, but almost universally recovered in both HFH-30364 and RN4850 strains. (52). Notably, mutations in *pgl* are known to affect cell-wall specific phenotypes, such as sensitivity to þ-lactam antibiotics and cell-wall permeable compounds (52, 53). In addition, *pgl* mutations increase the cell-surface charge of *S. aureus*, similar to our observations in evolved *ktr* mutants (53). This data suggests the potential for convergent evolution of spermine resistance at the mechanistic level, albeit via different genetic mechanisms. In the future, it would be interesting to investigate if strain-specific differences in cell-wall composition predispose spermine resistance to evolve via mutations in *ktr* or *pgl*.

While this study has identified new bacterial genes and pathways that contribute to polyamine resistance, the mechanism of polyamine toxicity in *S. aureus* remains mysterious. Previous efforts to characterize spermine toxicity revealed the gene *menD* as a potential target, revealing the electron carrier menadione as an important factor in mediating toxicity (16, 55). This study found that inactivation of *menD,* the gene that encodes for a menaquinone biosynthetic protein in *S. aureus,* increased resistance to spermine under aerobic conditions. While we did not recover mutations in *menD* during experimental evolution, it is possible that inactivating *menD* similarly alters the membrane potential and cell-surface charge of *S. aureus,* indicating a similar mechanism of resistance as discovered here (53, 55). However, it is more likely that the data in previous studies suggests that menadione interacts with polyamines and creates a toxic product that kills *S. aureus* (16). Ultimately, the mechanism of bacterial killing by polyamines will continue to be an important area of investigation and could additionally help in clarifying mechanisms of evolved polyamine resistance. Together this study reveals a new role for potassium transport in the unique polyamine susceptibility of *S. aureus*, with consequences for the evolution of multidrug resistance in this global human pathogen.

## Acknowledgements

We are grateful to members of the Barber lab and Tony Richardson for helpful discussions. This work was supported by National Institutes of Health grants to MFB (R35GM133652, R21AI173839). KC is a recipient of a National Institutes of Health Genetics Training Grant (T32GM149387). CHK is a recipient of the Helen Hay Whitney Foundation fellowship and L’Oréal USA FWIS fellowship. The authors declare no competing interests.

## Materials and Methods

### Strains

All strains used in the experimental evolution experiment were acquired from BEI resources. *Ktr* mutants were generated for the purposes of this study following an allelic exchange protocol described below.

### Experimental evolution

Glycerol stocks of each strain were struck out on to tryptic soy agar (BD) plates and incubated overnight at 37° C. For each strain, a total of 6 single colonies were picked, and each colony was used to inoculate a separate 3 mL culture containing tryptic soy broth (TSB) (BD). These liquid cultures were grown overnight for approximately 18 hours, shaking (225 rpm) at 37° C. Each liquid culture was used to initiate a replicate population in the evolution experiment. Four of the cultures were diluted 1/100 into fresh TSB containing 2mM of spermine (Sigma, 71-44-3), and two of the cultures were diluted 1/100 into plain TSB as a control. After passaging, the populations were returned to the shaking incubator. The populations were passaged daily in the same 1/100 dilution throughout the course of the experiment. Populations initially exposed to spermine on the first passaged were continuously exposed to increasing concentrations of spermine for the remainder of the evolution experiment. Control populations were passaged daily in plain TSB. Populations were sampled and frozen in a glycerol solution (25% final v/v) at passage 0, 4, 6, 8, and 10.

### DNA isolation and sequencing

Whole bacterial populations were sequenced that were exposed to spermine during the experiment, as well as the control populations that were never exposed to spermine. All four replicates of *S. aureus* MN8 populations were sequenced at passages 4, 6, 8 and 10. All four replicate populations of *S. aureus* HFH-30364 and *S. aureus* RN4850 that were exposed to spermine were sequenced at passage 10. For all experimentally evolved populations, ancestral clones for each strain that were used to initiate the evolution experiments were sequenced to provide a reference genome for variant calling. We also sequenced the two control populations (never exposed to spermine) for each strain background at passage 10. To isolate DNA, populations were struck out on to TSA plates from glycerol stocks, and mixed-colony samples were taken liberally from all parts of the plate to capture any diversity potentially present in the population. Bacterial samples were resuspended in TE buffer containing 50 µg/mL lysostaphin and incubated for one hour to facilitate lysis of cells. DNA was then harvested using a Qiagen DNeasy blood and tissue kit (CAT# 69504) according to manufacturer’s instructions. DNA extracts were sent to SeqCenter (seqcenter.com) for Illumina sequencing.

### Data processing and variant calling

Mutations in the evolved populations were identified with breseq v0.35.7 (58) using the default settings and polymorphism mode to calculate the frequencies of variants detected in the reads. Polymorphism mode in breseq calls a variant in the population if it is observed in both strands of at least 5% of the reads. The average read depth for MN8, HFH-30364, and RN4850 populations were 439x, 410x, and 600x respectively. Average genome coverage for MN8, HFH-30364 and RN4850 was 99.1%, 99.8%, and 99.5% respectively.

### Minimum inhibitory concentration measurements

Minimum inhibitory concentration (MIC) experiments were conducted using a modified broth microdilution method. To measure the MIC of a given compound, a single colony of bacteria was picked and inoculated into 4mL of TSB to start overnight cultures. The next day, the OD600 of the cultures was measured and diluted to an OD600 of 0.04. Different concentrations of an antimicrobial compound of interest (e.g. spermine) were made by diluting stocks of the chemical in 2X TSB and water, yielding the final concentrations of the compound in 1X TSB. 50 µL of diluted overnight culture was mixed 1:1 with each concentration of the prepared antimicrobial agent. Mixtures of bacterial cultures and antimicrobial compounds were incubated statically in 96 well plates at 37C for 24 hours. MIC values were determined as a range between the highest concentration at which visible growth occurred, and the lowest concentration at which no visible growth occurred. Bacteria were challenged at each concentration in triplicate, and the results reported are the average of three individual experiments.

### Protein modelling and mutation mapping

Published protein structures (PDB ID: 4J7C) and genetic sequences for the *Bacillus subtilis* KtrAB complex were used as guides to model the structure of the *S. aureus* KtrAD protein complex in ChimeraX (version 1.6.1).

### Generation of S. aureus mutants

To generate single base-pair (bp) mutations, allelic exchange was used with the pIMAY vector following protocols previously described (59). PCR products containing sequence approximately 1,000 bp upstream and downstream of the mutation of interest were generated. Primers were designed containing homology arms to facilitate assembly into the cloning vector. Genomic DNA extracted from the evolved MN8 isolates was used as template DNA to amplify the mutations of interest. PCR products were assembled into a digested pIMAY vector using NEBuilder HiFi DNA assembly (CAT# E2621S) according to the manufacturer’s instructions. Assembled vectors were chemically-transformed into *E. coli* DC10B and plated on to TSA + 25 µg/mL chloramphenicol to select for successful transformants. Colonies were screened with colony PCR to confirm presence of insert gene into pIMAY vector. Validated plasmids were isolated and transformed into recipient *S. aureus* strains as described previously (59). Allelic exchange of target genes were constructed according to Monk et al. 2012 (60). Transformed *S. aureus* cells were plated on to TSA + 10 µg/mL chloramphenicol at 28 C overnight. Single colonies were picked re-plated on to TSA + 10 µg/mL chloramphenicol to select for a single recombination event. Single colonies that grew on chloramphenicol were picked, grown up overnight in plain TSB, and then plated on to TSA + 1 µg/mL anhydrotetracycline (atc) to select for loss of the plasmid backbone. Successful secondary recombinants would be permissive to growth on atc. Single colonies that could grow on atc were picked and sequenced at the locus of interest to confirm that the mutation of interest was introduced successfully.

### Cytochrome C binding assay

Relative cell-surface charge was determined by measuring cytochrome C binding as described previously (61). Stationary phase overnight cultures were harvested and adjusted to an OD_600_ of ∼1.1 in 2mL of sodium acetate buffer (20 mM, pH 4.6). Cultures were washed twice in sodium acetate buffer before being resuspended in 0.5mL sodium acetate buffer with 0.25 mg/mL cytochrome C. Resuspended pellets were incubated while shaking at 37 °C for 15 minutes, then centrifuged at 1600 x g for 2 minutes. The supernatant was removed and aliquoted into 96 well plates, where the absorbance was measured at 410 nm using a Biotek plate reader. The percentage of cytochrome C bound for a given sample was calculated as the fraction of absorbance relative to 0.25 mg/mL cytochrome C in sodium acetate buffer alone.

### Chemically defined media lacking excess potassium

To measure the effect of specific amounts of potassium in the growth media, we constructed chemically defined media lacking excess potassium as described previously (37). Defined amounts of potassium were added into the media by the addition KCl or NaCl. Survival assays were conducted at different KCl or NaCl concentrations as described below.

### Survival assays

Overnight cultures were harvested, adjusted to a desired OD600 and incubated with an antibacterial compound of interest following the same protocol described above for MIC assays. Bacterial survival in various compounds was measured by plating and enumerating CFUs over the course of 24 hours of static incubation at 37 °C.

## Notes

### Competing Interest Statement

The authors have declared no competing interest.

