## Supplemental tables for "Evolution of polyamine resistance in *Staphylococcus aureus* through modulation of potassium transport"

| Passage | Spermine exposure (mM) | Passage saved? |
|---|---|---|
| 0 | 0 | yes |
| 1 | 2 mM | no |
| 2 | 2 mM | no |
| 3 | 2 mM | no |
| 4 | 4 mM | yes |
| 5 | 6 mM | no |
| 6 | 6 mM | yes |
| 7 | 6 mM | no |
| 8 | 6 mM | yes |
| 9 | 7 mM | no |
| 10 | 7 mM | yes |

**Supplemental table 1:** Spermine concentrations that experimentally evolved populations were exposed to at each passage of the evolution experiment*.* Spermine was added directly to the growth media resulting in the final concentration displayed in the table.

| Strain | Source | Details |
|---|---|---|
| *S. aureus* MN8 | BEI resources | https://www.beiresources.org/Catalog/bacteria/HM-162.aspx |
| *S. aureus* HFH-30364 | BEI resources | https://beiresources.org/Catalog/bacteria/NR-10189.aspx |
| *S. aureus* RN4850 | BEI resources | https://www.beiresources.org/Catalog/bacteria/NR-45955.aspx |
| *S. aureus* MN8 KtrA*:*H47Y | This study | Tyrosine substituted for histidine at the $47^{th}$ position of the KtrA protein in the MN8 strain background |
| *S. aureus* MN8 KtrA*:*G94V | This study | Valine substituted for glycine at the $94th^{th}$ position of the KtrA protein in the MN8 strain background |
| *S. aureus* MN8 KtrA*:*H47Y-G94V | This study | Tyrosine substituted for histidine at the $47^{th}$ position of the KtrA protein and valine substituted for glycine at the $94th^{th}$ position of the KtrA protein in the MN8 strain background |

**Supplemental table 2:** List of S. aureus strains and their sources used in this study*.* Single amino acid mutations were introduced into the MN8 parent strain by allelic exchange as outlined in the methods.

| Gene | Allele | Strain | Metadata | Sequence ID |
|---|---|---|---|---|
| *ktrA* | **A178V** | n/a | clinical or host-associated sample of *S. aureus* | HDB4168921.1 |
| *ktrA* | A178T | SA-71 | n/a | HDB2454818.1 |
| *ktrA* | E151V | CFSA200 | clinical or host-associated sample from *S. aureus* | WP_086896505.1 |
| *ktrA* | E151K | n/a | clinical or host-associated sample from *S. aureus* | HDB5855606.1 |
| *ktrA* | E76K | SMC9559 | Generic sample Host: *Homo sapiens* | WP_217801415.1 |
| *ktrA* | | AF254 | clinical or host-associated sample of *S. aureus* | MBV2625534.1 |
| *ktrA* | **H47Y** | C51 | Pig isolate | WP_031790491.1 |
| *ktrA* | | DAR3583 | Generic sample Host: *Homo sapiens* | WP_031790491.1 |
| *ktrA* | | RIVM_M084907 | Pathogen: clinical or host-associated sample from Staphylococcus aureus | HDE6238165.1 |
| *ktrA* | | RIVM_M085009 | Pathogen: clinical or host-associated sample from Staphylococcus aureus | HDE6260146.1 |
| *ktrA* | | RIVM_M043274 | Pathogen: clinical or host-associated sample from Staphylococcus aureus | HDF4313963.1 |
| *ktrD* | **G94V** | H88163 | Generic sample Host: *Homo sapiens* | WP_031807927.1 |
| *ktrD* | G94D | M8158 | Pathogen: clinical or host-associated sample from Staphylococcus aureus | HDA5546730.1 |
| *ktrD* | **Q450K** | CM60 | Pathogen: clinical or host-associated sample from Staphylococcus aureus | WP_111088627.1 |

**Supplemental table 3:** Mutations in *ktrA* and *ktrD* genes in publicly available sequences of *S. aureus* isolates. Queries were conducted using the identical protein group database on NCBI.

| Replicate Line | *ktrA* | *ktrD* |
|---|---|---|
| 1 | H47Y | G94V |
| 2 | - | - |
| 3 | E76G | M391I |
| Control | - | - |

**Supplemental table 4:** Mutations that arose in *S. aureus* MN8 *ktrA* and *ktrD* during a pilot evolutionary experiment.
